# Runaway resorption of microcracks contributes to age-related hip-fracture patients

**DOI:** 10.1101/2024.06.09.598123

**Authors:** Marena Gray, Oliver Boughton, Crispin Wiles, Christina Reinhard, Nghia T. Vo, Robert Atwood, Richard Stavri, Justin P. Cobb, Ulrich Hansen, Richard L. Abel

## Abstract

Microdefects, including microcracks and resorption trenches, may be important contributors to bone fragility. 3D microdefect morphology was imaged using synchrotron micro-CT to develop a classification system for investigating the relationship with bone mechanics and hip-fractures.

Femoral heads from ageing hip-fracture patients (*n*=5, 74-82 years) were compared to ageing non-fracture controls (*n*=5, 72-84 years). Two trabecular cores were prepared from the chiasma; one was imaged using synchrotron micro-CT to measure microdefects and one was mechanically tested to measure tensile strength. Morphological and mechanical data were compared and correlated using Mann Whitney U test and Pearson’s rank correlation.

Microdefects varied and were classified into four categories based on shape and measurable parameters. Hip-fracture donors exhibited significantly higher density of all microdefects (*p*<0.05). Microdefect volume was strongly negatively correlated with ultimate tensile strength (*p*<0.05) and stiffness (*p*<0.05).

Microdefects might contribute to loss of bone strength and fragility fracture via runaway resorption. Microcracks could promote focussed osteoclastic resorption and the formation of resorption pits which create stress risers leading to the re-formation of microcracks under continued load. CT-based classification methods should be used to explore the complex interaction between microdefects, metabolism, and bone fracture mechanics.

## Introduction

The role of 3D trabecular microarchitecture in bone mechanics and fragility has been extensively analysed^2–6,9^ but 3D microdefects, which could undermine architecture, have been less well studied^6,7^. Microdefects are damage that has been formed at the microscopic level of bone^8–10^, in the bone matrix^9^, and has been determined to be a normal physiological response to skeletal loading^11–15^. Microdefects have traditionally been assessed using 2D histomorphometry methods^1,9,15–18^ which do not reveal the 3D shape of defects for identification^7,19^. Benchtop micro-computed tomography (micro-CT) systems have also been used but specimens require radio-opaque staining which may not show all of the microcracks, and resolution is limited to tens of μm resolution^7,20^. Recent developments in Synchrotron Micro-CT allow for rapid scanning (approximately five to twenty minutes) at micron-level resolution ^7^. As such, it may be possible to investigate 3D microdefect morphology for biomedical research and address key questions about the role of microdefects in bone mechanics, health, and disease.

### Microdefect types

Histomorphometry studies have identified three types of microdefects which are likely interrelated and form together. Microcracks are defined as fractures in bone tissue at the scale of 50-100μm^9,21^. Diffuse damage consists of networks of small sub-lamellar size cracks (approximately 1 μm)^22^. Osteoclastic resorption pits^23–25^ usually appear as concavities in the bone surface or cylindrical tunnels in the tissue and are approximately 8-16μm^25^ long. Within the bone matrix itself, there are many physiological discontinuities, not limited to vascular canals, canaliculi and osteocyte lacunae. The latter have been the subject of interest, also under the scrutiny of Micro-CT imaging^26^ Microcracks are a known stimulus for osteoclastic remodelling^1,11–13,16,23,27,28^ and microcracks undergoing targeted resorption have been observed in histology sections^29^. Osteoclastic resorption may also create stress risers that act as a stimulus and nucleation point for more cracks^24,30^. Therefore, the formation of microcracks and resorption pits are related and should be studied together.

### Microdefects and Mimechanical properties

All three types of microdefects could undermine the microstructure of a bone and reduce the mechanical properties increasing the risk of a fracture during a bump or fall. Therefore, it is important to determine whether microcracks and resorption pits occur together in bone tissue or in isolation. The effect of microdefects on mechanical properties and fracture mechanics is likely complex, especially given that the two are interrelated because microcracks are resorbed by osteoclasts. Ultimately all fractures of the cortical and trabecular bone must begin as microscopic or sub-microscopic linear microcracks^31,32^ which propagate into a fracture, usually under a traumatic load^8,18^. However, microcracks are also thought to be a key component in dissipating energy and maintaining the mechanical properties of bone^3,4,8,9,33–36^. Therefore, microcrack formation is also considered a normal physiological process which reduces fracture risk by absorbing energy from fatigue or physiological loading^5,11,12,20,37^. When microcracks are removed and replaced with new tissue by the remodelling process there should be no long-lasting impairment of the tissue.

### Classifying and quantifying microdefects morphology

Studies investigating the relationship between bone fracture mechanics, fragility and crack morphology are required to determine whether microcracks, osteoclastic resorption pits, or both, are an important contributor to bone strength or fragility. The morphology of microdefects is complex and there is a scarcity of data on size, shape, and density. Two classifications of microdefects have been explored based on location within the bone^10,18^, and the number or density^9,35,38,39^ of microdefects. For example, Norman and Wang^18^ classified microdefects based on location relative to cement lines in both human femora and tibiae. This method of classification is useful with regards to the presence of linear microcracks in certain sections of the bone and the role of osteons but is not useful in exploring relationships between different types of microdefects and bone integrity. The criteria defined for linear microcracks and diffuse damage was limited to comparisons with other structures, for example linear microcracks were smaller than vascular channels but larger than the canaliculi^10,38^. Previous classification systems of microdefects relied on 2D images or computer simulations^2,10,23,36^, which have been unable to distinguish partially resorbed microdefects to the detail seen now. Previous simulations and models, e.g. the lamellar wood model, are oversimplified and therefore are not representative of how microdefects form^5,40^. Therefore, a system for classifying and quantifying microdefects is necessary to investigate the role of microcracks and resorption pits in bone health and disease.

### Imaging microdefects

The use of synchrotron-CT is a relatively new modality in the imaging of bone, which has already demonstrated its effectiveness in imaging microcracks^6,7,33,41^ but limited access to the technology has meant 2D histology images and 3D benchtop CT scans remain the most common methods of analysis. The advantage of synchrotron CT is that the combination of higher resolution and 3D imaging can be applied to image, classify, and measure microdefects and fully differentiate microcracks at different stages of remodelling and resorption pits. A few studies have already used the technique to image microdefects but there was no classification system for differentiating according to the shape and scale^6,7,19,33,41^.

### Aims and objectives

The aim of this work was to investigate 3D microdefect morphology in human bone tissue from hip fracture patients versus ageing non-fracture controls: including microcracks and osteoclastic resorption pits. There were three objectives: i) image microdefects using synchrotron X-ray micro-CT and investigate the range of variation of microdefect shape and scale, ii) develop a classification system and iii) test the application by comparing microdefects in trabecular bone from hip-fracture patients with non-fracture controls and correlating microdefects with mechanical strength and stiffness.

## Materials and Methods

### Donors and sample preparation

Trabecular bone cores were prepared from the femoral heads of two groups of patients: a hip-fracture group and non-fracture control donors^19,41^. Neither group had been treated with bisphosphonates or other bone metabolic treatments prior to the study. The fracture group was sourced from patients who had hip arthroplasty surgery for fractures of the femoral neck within the Imperial College Healthcare National Service Trust in London. The samples from the non-fracture control group were taken from cadavers. The exclusion criteria included those with a history of primary or secondary bone disease. All the procedures performed were in accordance with the ethical standards of the Imperial College Tissue Bank (R13004) and the 1984 Declaration of Helsinki. In total, eight samples were taken from hip-fracture patients and five from non-fracture control donors. These samples were 10mm in height and 7mm in diameter and were taken from the primary compressive trabecular arcade of the femoral heads. The cores were stored at -80 C, and remained in the storage facility unless directly being used for testing,

### Synchrotron X-ray micro-CT

Imaging was carried out at Diamond Light Source, United Kingdom, with Synchrotron X-ray micro-CT using Beamline I12^42^ on only five cores from fracture patients and five cores from non-fracture controls, due to limited experiment time with the synchrotron. The setting parameters used were photon energy 53 keV, image volume 23.3⍰mm^3^, voxel resolution 1.38⍰mmμm^3^/voxel, 6400 projections, 180° rotation. Images were constructed in-house using GPU enabled Filtered Back Projection within the DAWN^41^ package, and using Titarenko’s ring suppression method^43^ and then linked to image analysis software ImageJ and VG Studio Max^44^ to allow for identification of microdamage.

### Microdefect assessment

From a given bone sample, a central fiducial volume of dimensions 3.28mm in diameter and 2.76mm in height was taken, to reduce the likelihood of introducing artefacts from the cutting and drilling process. The volume of interest was segmented with an automated global threshold technique that was applied to identify the trabecular bone tissue (including the microdefects)^6,45^. Bone volume fraction (BVF) was calculated by dividing the total number of voxels that represented bone by the total number of voxels in the whole scan volume^41,44^. Subsequently, the microdefects within the bone were visually identified and segmented by masking the microcracks in VG Studio Max 2.2 (Volume Graphics)^42^ then inspected to qualify the morphology and measured quantitatively. There were, in total, 2000 continuous slices individually examined in three planes. Microdefect density per mm^3^ was calculated by normalising the number of defects to BVF. Microdefect volume was measured counting the segmented voxels. Microdefect length was established as the longest axis and measured by calculating the length of the bounding cuboid of the microdefect orientated in the primary axis. Microdefect volume was calculated by converting the number of voxels into mm^3^ (1 voxel being the equivalent of 1.3µm^3^).

### Two step classification system for microdefects

In the process of developing a classification system for microdefects based on shape and size, it became clear there were four types visible in synchrotron micro-CT images, including resorption pits, microcracks, resorption trenches and partially resorbed microcracks (Fig. 1). The presence of partially resorbed microcracks visible in the synchrotron scans demonstrate that existing classification systems^10^ for microdefects do not consider the full range of variation in type. Therefore, a two-step classification system was developed (Fig. 1) which incorporated the full range of variation in microdefects imaged. The first step involves categorising predominately by shape, which can be better appreciated in 3D, into three different groups. Parameters of length, width, depth, and volume were then used in the second step to differentiate within a class. These parameters are based on numerous analyses in the literature, with microcracks noted to be a length of 50-100µm^4,9,21^ and resorption pits measured at 8-16µm long^23,30^. This process is outlined in Table 1 and Figure 1.

**Table 1:**
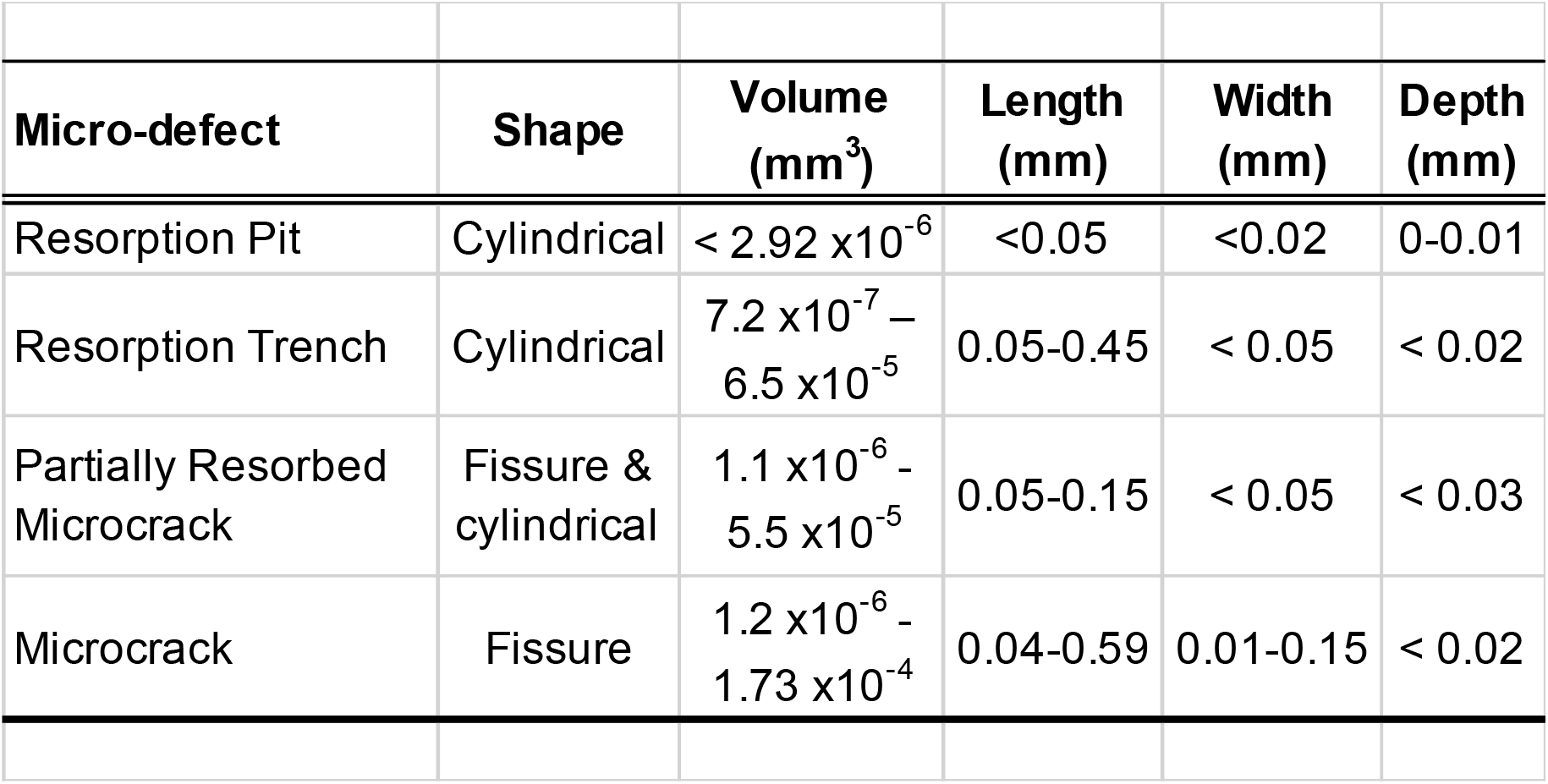
Classification of each microdefect was categorised by the volume, length, width, depth and the shape of each element.

**Figure 1:**
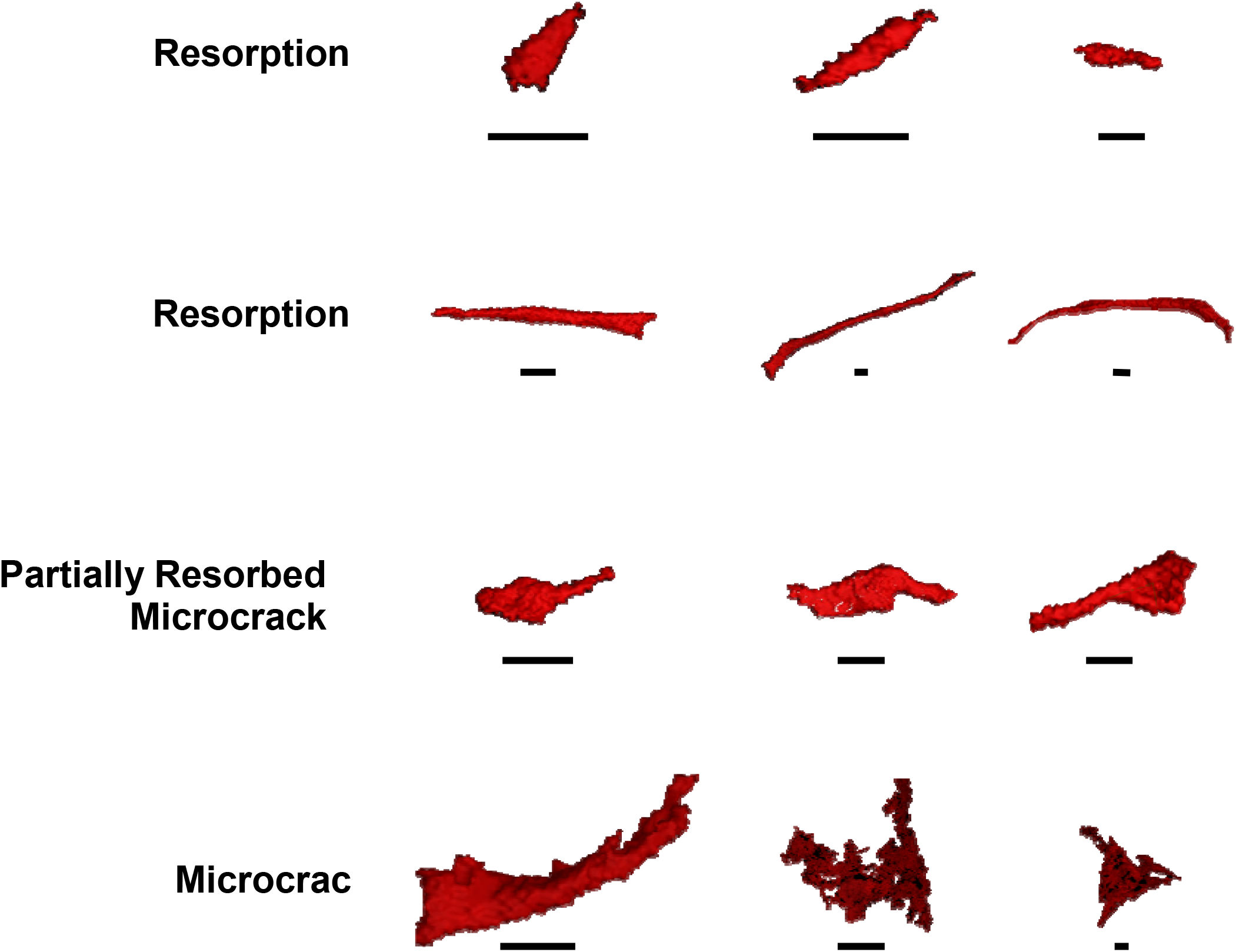
Examples of the four categories of microdefects classified: resorption pits, resorption trenches, partially resorbed microcracks and microcracks. Scale bars measure 0.02mm.

### Mechanical testing

Ten rectangular-shaped testing samples were harvested immediately adjacent to the region of the cylindrical cores7,41. The specimens were 11⍰ mm in height, 2.8⍰ mm in width, and 1⍰ mm in depth. The ends of each sample were potted in bone cement, which served as clamps and stored at −80° ⍰ C until testing. During tensile testing, the samples were kept hydrated in the fluid chamber built into the micromechanical device7,41. All specimens were static then load-controlled tensile tested with a fixed strain rate at 0.001⍰ s−1 using a custom-built micromechanical tensile test rig, originally designed for previous projects7,41. The tests were conducted at room temperature. After loading, the stress-strain curves were examined to identify the Young’s modulus and ultimate tensile strength (UTS), which were normalised according to trabecular bone volume.

### Statistics

Statistical analyses were performed using IBM SPSS Statistics 23 (Armonk, New York) and the graphs were generated with GraphPad Prism 8 (San Diego, California). Donor groups were compared using parametric descriptive statistics with 2Way Anova. Microstructure and tissue mechanics were correlated using Pearson’s correlation co-efficient.

## Results

Synchrotron images successfully visualised and reconstructed a wide range of microdefects with varied morphology (Fig. 1). The morphology of the microdefects was consistent with linear microcracks, partially resorbed microcracks and osteoclastic resorption pits but not diffuse damage.

The mean age of the non-fracture controls and hip fracture-donors was similar (p>0.05, Table 2). Compared to the controls, the hip-fracture donor group exhibited a significantly greater density of microcracks, partially resorbed microcracks and resorption trenches (p<0.05, Table 2).

**Table 2:** Donor demographics and data. Means and (StDev) compared using unpaired t-test. There was a significant increase in the density of resorption trenches, microcracks and partially resorbed microcracks in the fracture group, as well as a significant reduction in UTS in the fracture group. Young’s modulus was similar between the fracture group and the control.

|  | Non-Fracture Controls | Hip-Fracture | *p* |
| --- | --- | --- | --- |
| Mean age, yrs (SD) | 78.6 (5.6) | 77.8 (3.4) | 0.775 |
| Female, proportion (%) | 4 (80.0) | 5 (100.0) | n/a |
| Male, proportion (%) | 1 (20.0) | 0 (0.0) | n/a |
| Resorption pit (density/mm<sup>3</sup>) | 0.281 | 0.283 | 0.774 |
| Resorption trench (density/mm<sup>3</sup>) | 0.121 | 2.097 | <0.001 |
| Partially resorbed cracks (density/mm<sup>3</sup>) | 0.241 | 1.190 | <0.001 |
| Microcrack (density/mm<sup>3</sup>) | 0.482 | 3.458 | <0.001 |
| Ultimate Tensile Strength, MPa (SD) | 49.22 (5.73) | 30.80 (4.59) | <0.001 |
| Young's Modulus, N/m<sup>2</sup> (SD) | 7.51 (2.60) | 5.19 (1.03) | 0.146 |

The density of resorption pits was similar in the non-fracture controls and hip fracture donors (p>0.05, Table 2).

Bone strength and stiffness were negatively correlated with the average volume of microdefect in the bone sample. Normalised UTS was strongly and negatively correlated with the average volume of microdefects (-r^2^ = 0.82, *p*<0.05, Fig. 2), which is defined as the total volume of microdefects in a bone core divided by the frequency of microdefects in a bone core sample. Normalised Young’s modulus was negatively correlated with the average volume of microdefects (-r^2^ = 0.53, *p*>0.05, Fig. 2) but was not significant.

**Figure 2:**
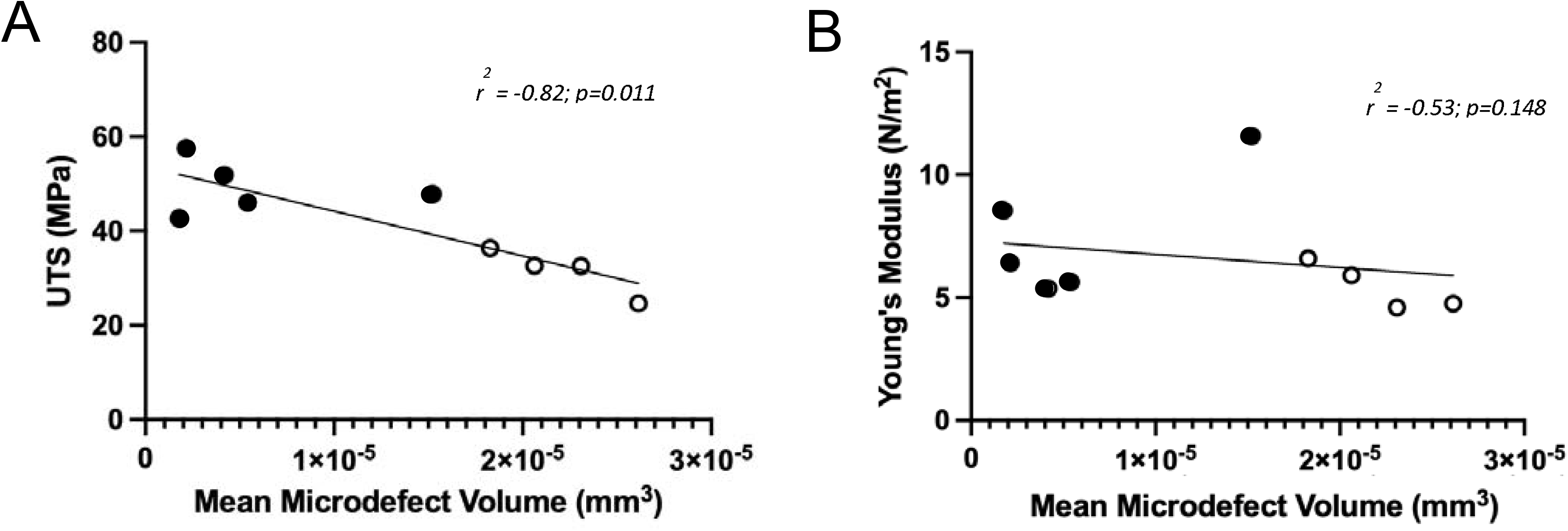
Bone strength and stiffness are negatively correlated with the mean volume of defect in the bone sample. A) UTS is average volume of microscopic bone defects (*r*^2^ = 0.82, p<0.05). B) Young’s Modulus is negatively correlated with the average volume of microscopic bone defects (*r*^2^ = 0.53, *p*>0.05). Spearman’s r-correlation with Welch’s t-test.

Negative regressions were calculated for each category of microdefect across both ageing fracture and non-fracture groups compared to average UTS (Fig. 3). There was a significant negative regression between average UTS and microcracks (p<0.05, r^2^ = 0.2915, Fig. 3). The average UTS compared to partially resorbed microcracks (p>0.05, r^2^ = 0.1496), resorption pits (p>0.05, r^2^ = 0.00085), and resorption trenches (p>0.05, r^2^ = 0.1363) were found to not have a significant negative regression.

**Figure 3:**
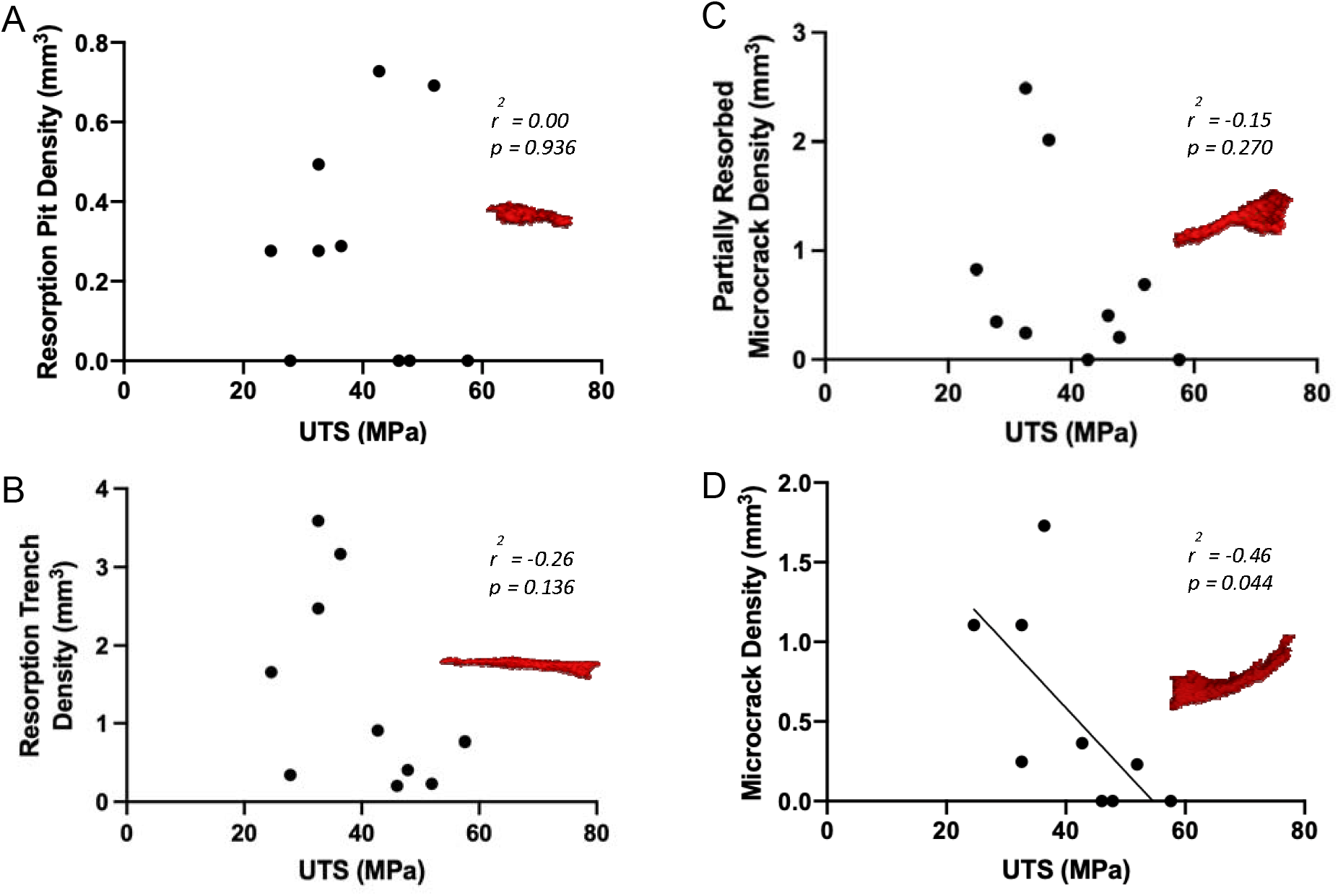
Bone ultimate tensile strength has a negative regression with the average density of defect per volume of the bone sample in all types of microdefects. Examples of each are given. A) Resorption Pit, B) Resorption Trench, C) Partially Resorbed Microcrack, D)

## Discussion

The aim was to investigate 3D microdefect morphology in human bone tissue by comparing data between hip fracture donors versus ageing non-fracture controls. 3D images of microdefects were successfully obtained using synchrotron micro-CT scans revealing wide variation in morphology which could be categories into four types: microcracks, resorption pits, resorption trenches and partially resorbed microcracks. Microcracks and resorption trenches are similar in size but have divergent shapes: microcracks are fissures whilst resorption trenches are cylindrical (Fig. 1). Resorption trenches are approximately ten times longer than resorption pits, although both are cylindrical in shape (Fig. 1).

Hip-fracture donors exhibited a significantly higher density of microcracks, partially resorbed microcracks and resorption trenches compared to ageing controls (Table 2). Further, hip-fracture patients exhibited significantly lower tensile strength and showed lower stiffness (Fig. 2). Inferior mechanics of hip-fracture patients could be attributed to the increased density and volume of defects, which reduce the connectivity of the elements and the cross-sectional area for resisting loads, ultimately leading to failure of the trabecular architecture. The presence of a high proportion of partially resorbed microcracks in hip-fracture patients relative to controls suggests osteoclastic resorption of microcracks could be a cellular mechanism contributing to reduced strength, therefore contributing to age-related fragility fractures at the hip.

### Age-related fractures are associated with accumulated microdefects

Synchrotron microdefect data support the theory that microdefects contribute to a reduction in trabecular bone strength and increase the risk of a hip-fracture after a bump or fall^7,19,41^. Microcrack density was five-fold greater in the fracture group and resorption trench density was twelve-fold in the fracture group compared to non-fracture controls. There was no significant difference in the density of resorption pits between either group. There was a significant negative correlation in bone strength compared to average microdefect volume in bone tissue (-r^2^ = 0.82, *p*<0.05, Fig. 2). There was also a negative correlation in bone stiffness compared to average microdefect volume in bone tissue (-r^2^ = 0.53, *p*>0.05, Fig. 2), however this was not significant. This suggests stiffness is a mechanical property representative of the structure as a whole, hence why stiffness is typically affected by total mineral density and metabolic disease states^15,47,48^ and is only moderately reduced^49^ in this analysis. Comparatively, ultimate tensile strength is representative of the structure at a specific tested point, therefore an accumulation of microdefects could exceed the intrinsic repair capability of the bone, resulting in further damage and potentially failure of the bone, therefore strongly reduce bone material strength.

These conclusions support and are supported by the literature. Prendergast and Taylor suggest there is the presence of homeostatic microdefects and remodelling only occurs if a critical value is reached^27^. In a review paper, Seref-Ferlengez discussed how when bone tissue structure becomes compromised, microdefect accumulation is a significant contributor to future fracture risk^5,40^. Zioupos and colleagues determined that changes in mechanical properties were caused by microdefect occurrence, rather than microdefects being an artefactual consequence^4^. Yeh and Keaveny explored loading trabecular bone and found that extensive microdefects were primarily responsible for the loss in strength and stiffness^50^.

### Interaction between resorption pits and microcracks causes runaway resorption

The synchrotron micro-CT images illustrate how microdefects exist on a spectrum and likely reflect the interaction in formation. For example, when a microcrack undergoes targeted remodelling by osteoclasts, this was frequently visible as microcracks undergoing active resorption by osteoclasts (Fig. 1). Burr and Allen initially proposed a theory that an age-associated failed feedback loop involving remodelling in response to cracks may precipitate microdefect accumulation (both microcracks and resorption pits), which reduce the strength and stiffness (see Fig. 2) and so promote bone fractures^9,11,13,14^.

When a microcrack forms as consequence of applied stress, generating tissue strain^5^, a targeted process of osteoclastic resorption would remove the crack. However, in the absence of osteoblastic bone formation, the cavity cannot be filled. Any unfilled resorption pits increase the area of stress concentration, leading to further microcracks, resorption and extension of the pits into a resorption trench. At a certain depth^51^, runaway resorption^2,23,24,30,51^ fundamentally undermines the architecture of the tissue^2,51^, compromising strength. This relationship has been addressed using computer simulations^24,51^, where in-silico models suggest that resorption trenches are formed from resorption of pits due to continuous remodelling of the same area of tissue, rather than a single resorption event^24^. The authors noted that pits greater than 32μm in depth were associated with resorption trench formation because osteoblasts were not able to fill such a large space. Experimental and modelling studies are required in combination to create a platform for investigating the aetiology of fragility fractures with a classification system. Slyfield et al. demonstrated significant associations between the presence of microdefects and resorption pits, which is supported by the regression data in Figure 3. As the only significant contributor to a significant reduction in ultimate tensile strength was the microcrack subcategory, this suggests that microcracks are the more pathological type of microdefect^5,16^.

This study acknowledges there may be several causes of osteoclastic resorption trench formation. With the variability in the dimensions of microcracks and resorption trenches (Table 1), the presence of a resorption trench could also be due to the resorption of a large microcrack, rather than solely a consequence of concentrated stress in the tissue. At this stage, it is difficult to differentiate between the two aetiologies. This study also acknowledges the need for expanding this work further to include greater numbers of samples from individuals of varying demographics to allow for more refinement of the classification system to ensure its reproducibility.

Microdefects are likely a major component of bone fragility and hip-fractures that require further study. The microdefect research highlights the importance of risk factors other than bone mineral density in predicting bone strength and fracture risk. The increased density of microcracks and the phenomena of runaway resorption of osteoclastic resorption pits could increase the potential risk for an age-related fragility fracture during a trip or fall.

## Conclusion

This paper explored the morphology of microcracks and osteoclastic resorption pits in human bone tissue. Synchrotron micro-CT imaging successfully captured microdefects and distinguished microcracks, resorption pits and partially resorbed microcracks using a two-step classification system. The classification system presented in this paper has the potential to create a platform to study the role of microscopic defects in whole bone mechanics. Bone tissue samples from fracture patients exhibited significant greater densities of microcracks and resorption pits in hip-fracture patients than in controls. There is a likely interaction between microcracks and osteoclastic resorption pits, which may be integral to the development of fragility fractures. Microcracks likely act as a stimulus for resorption which can predispose to the generation of either physiological osteoclastic resorption pits and successful bone formation or runaway resorption leading to ultimate failure of bone. Future studies of the relationship between hierarchal bone structure and fracture mechanics need to take microdefects into account.

## Acknowledgements

We wish to thank the direct care teams at St Mary’s Hospital for consenting patients and collecting tissue samples. The authors would also like to thank the patients who consented to donate tissue for research and the Imperial College Healthcare National Health Service (NHS) staff and Imperial Tissue Bank staff who helped with the collection of the samples. The research was funded by the Wellcome Trust and Engineering and Physical Sciences Research Council (EPSRC) Osteoarthritis Centre of Excellence (088844/Z/09/Z), the Michael Uren Foundation and the Science and Technology Facilities Council (STFC) Impact Acceleration Grant. The Diamond Light Source funded beam time allocations (EE0852, EE9811, SM10458, EE11204, SM13337). The authors are very grateful for the help of the Diamond Light Source staff at Harwell, UK.

## The authors declare no competing financial interests

### Author Contributions Study Design

JPC, UH and RLA. Study conduct: MG. Data Collection: MG, OB, CR, NTV, RA. Data Analysis: MG, UH, RLA. Data Interpretation: MG, UH, CW, RLA. Drafting Manuscript: MG, JPC, UH, CW, RS, RLA. Approving final version of manuscript: All. MG takes responsibility for the integrity of the data analysis.

## Data Availability Statement

Raw data can be made available to other authors on request.

## References

1. O’Brien, F. J., Brennan, O., Kennedy, O. D. & Lee, T. C. Microcracks in cortical bone: how do they affect bone biology? Current osteoporosis reports (2005). doi:10.1007/s11914-005-0002-1

2. Van Oers, R. F. M., Van Rietbergen, B., Ito, K., Huiskes, R. & Hilbers, P. A. J. Simulations of trabecular remodeling and fatigue: Is remodeling helpful or harmful? Bone (2011). doi:10.1016/j.bone.2011.01.011

3. Zioupos, P. & Currey, J. D. The extent of microcracking and the morphology of microcracks in damaged bone. J. Mater. Sci. (1994). doi:10.1007/BF00351420

4. Zioupos, P., Hansen, U. & Currey, J. D. Microcracking damage and the fracture process in relation to strain rate in human cortical bone tensile failure. J. Biomech. (2008). doi:10.1016/j.jbiomech.2008.07.025

5. Seref-Ferlengez, Z., Kennedy, O. D. & Schaffler, M. B. Bone microdamage, remodeling and bone fragility: how much damage is too much damage? Bonekey Rep. (2015). doi:10.1038/bonekey.2015.11

6. Larrue, A. et al. Synchrotron radiation micro-CT at the Micrometer scale for the analysis of the three-dimensional morphology of microcracks in human trabecular bone. PLoS One (2011). doi:10.1371/journal.pone.0021297

7. Ma, S. et al. Synchrotron Imaging Assessment of Bone Quality. Clinical Reviews in Bone and Mineral Metabolism (2016). doi:10.1007/s12018-016-9223-3

8. Burr, D. B. Why bones bend but don’t break. J. Musculoskelet. Neuronal Interact. (2011).

9. Burr, D. B. et al. Bone microdamage and skeletal fragility in osteoporotic and stress fractures. Journal of Bone and Mineral Research (1997). doi:10.1359/jbmr.1997.12.1.6

10. Dominguez, V. M. & Agnew, A. M. Microdamage as a Bone Quality Component: Practical Guidelines for the Two-Dimensional Analysis of Linear Microcracks in Human Cortical Bone. JBMR Plus (2019). doi:10.1002/jbm4.10203

11. Allen, M. R. & Burr, D. B. Bone Growth, Modeling, and Remodeling. in Basic and Applied Bone Biology (2019). doi:10.1016/b978-0-12-813259-3.00005-1

12. Lee, T. C., Staines, A. & Taylor, D. Bone adaptation to load: Microdamage as a stimulus for bone remodelling. J. Anat. (2002). doi:10.1046/j.1469-7580.2002.00123.x

13. Allen, M. R. & Burr, D. B. Skeletal microdamage: Less about biomechanics and more about remodeling. Clin. Rev. Bone Miner. Metab. (2008). doi:10.1007/s12018-008-9015-5

14. Allen, M. R., Burr, D. B. & Turner, C. H. The Effect of Age on Material Properties. in Osteoporosis in Men (2010). doi:10.1016/B978-0-12-374602-3.00018-3

15. Pagano, S. L. UKnowledge The Effect of Varying Bisphosphonate Treatment on Changes in Bone Microdamage in Osteoporotic Women. (2016).

16. Brandi, M. L. Microarchitecture, the key to bone quality. Rheumatol. (United Kingdom) (2009). doi:10.1093/rheumatology/kep273

17. Donahue, S. W. & Galley, S. A. Microdamage in bone: Implications for fracture, repair, remodeling, and adaptation. Critical Reviews in Biomedical Engineering (2006). doi:10.1615/CritRevBiomedEng.v34.i3.20

18. Wang, Z. & Norman, T. L. Microdamage of human cortical bone: Incidence and morphology in long bones. Bone (1997). doi:10.1016/S8756-3282(97)00004-5

19. Jin, A. et al. The effect of long-term bisphosphonate therapy on trabecular bone strength and microcrack density. Bone Jt. Res. (2017). doi:10.1302/2046-3758.610.BJR-2016-0321.R1

20. Wang, X. & Niebur, G. L. Microdamage propagation in trabecular bone due to changes in loading mode. J. Biomech. (2006). doi:10.1016/j.jbiomech.2005.02.007

21. Burr, D. B. The complex relationship between bone remodeling and the physical and material properties of bone. Osteoporosis International (2015). doi:10.1007/s00198-014-2970-4

22. Alvandi, L. M. Diffuse Damage Repair Mechanism in Bone. (2019).

23. Mcnamara, L. M. & Prendergast, P. J. Perforation of cancellous bone trabeculae by damage-stimulated remodelling at resorption pits: A computational analysis. in European Journal of Morphology (2005). doi:10.1080/09243860500096289

24. Mulvihill, B. M., McNamara, L. M. & Prendergast, P. J. Loss of trabeculae by mechano-biological means may explain rapid bone loss in osteoporosis. J. R. Soc. Interface (2008). doi:10.1098/rsif.2007.1341

25. Merrild, D. M. H. et al. Pit- and trench-forming osteoclasts: A distinction that matters. Bone Res. (2015). doi:10.1038/boneres.2015.32

26. Dong, P. et al. 3D osteocyte lacunar morphometric properties and distributions in human femoral cortical bone using synchrotron radiation micro-CT images. Bone 60, 172–185 (2014).

27. Prendergast, P. J. & Taylor, D. Prediction of bone adaptation using damage accumulation. J. Biomech. (1994). doi:10.1016/0021-9290(94)90223-2

28. Herman, B. C., Cardoso, L., Majeska, R. J., Jepsen, K. J. & Schaffler, M. B. Activation of bone remodeling after fatigue: Differential response to linear microcracks and diffuse damage. Bone (2010). doi:10.1016/j.bone.2010.07.006

29. Warden, S. J., Davis, I. S. & Fredericson, M. Management and prevention of bone stress injuries in long-distance runners. Journal of Orthopaedic and Sports Physical Therapy (2014). doi:10.2519/jospt.2014.5334

30. McNamara, L. M., Van Der Linden, J. C., Weinans, H. & Prendergast, P. J. Stress-concentrating effect of resorption lacunae in trabecular bone. in Journal of Biomechanics (2006). doi:10.1016/j.jbiomech.2004.12.027

31. Reeve, J. Role of cortical bone in hip fracture. Bonekey Rep. (2017). doi:10.1038/bonekey.2016.82

32. O’Brien, F. J. et al. The behaviour of microcracks in compact bone. in European Journal of Morphology (2005). doi:10.1080/09243860500096131

33. Zimmermann, E. A. et al. Age-related changes in the plasticity and toughness of human cortical bone at multiple length scales. Proc. Natl. Acad. Sci. U. S. A. (2011). doi:10.1073/pnas.1107966108

34. Ritchie, R. O. How does human bone resist fracture? in Annals of the New York Academy of Sciences (2010). doi:10.1111/j.1749-6632.2009.05232.x

35. Burr, D. B. Repair mechanisms for microdamage in bone. Journal of Bone and Mineral Research (2015). doi:10.1002/jbmr.2441

36. Mohsin, S., O’brien, F. J. & Lee, T. C. Microcracks in compact bone: A three-dimensional view. J. Anat. (2006). doi:10.1111/j.1469-7580.2006.00554.x

37. Lee, T. C. et al. Detecting microdamage in bone. Journal of Anatomy (2003). doi:10.1046/j.1469-7580.2003.00211.x

38. Burr, D. B. & Stafford, T. Validity of the bulk-staining technique to separate artifactual from in vivo bone microdamage. Clin. Orthop. Relat. Res. (1990). doi:10.1097/00003086-199011000-00047

39. Diab, T. & Vashishth, D. Effects of damage morphology on cortical bone fragility. Bone (2005). doi:10.1016/j.bone.2005.03.014

40. Seref-Ferlengez, Z., Basta-Pljakic, J., Kennedy, O. D., Philemon, C. J. & Schaffler, M. B. Structural and mechanical repair of diffuse damage in cortical bone in vivo. J. Bone Miner. Res. (2014). doi:10.1002/jbmr.2309

41. Ma, S. et al. Long-term effects of bisphosphonate therapy: Perforations, microcracks and mechanical properties. Sci. Rep. (2017). doi:10.1038/srep43399

42. Drakopoulos, M. et al. I12: The Joint Engineering, Environment and Processing (JEEP) beamline at Diamond Light Source. J. Synchrotron Radiat. (2015). doi:10.1107/S1600577515003513

43. Titarenko, S., Withers, P. J. & Yagola, A. An analytical formula for ring artefact suppression in X-ray tomography. Appl. Math. Lett. 23, 1489–1495 (2010).

44. Abel, R. L., Laurini, C. R. & Richter, M. A palaeobiologist’s guide to ‘virtual’ micro-CT preparation. Palaeontol. Electron. 15, (2012).

45. Yan, L. et al. A method for fracture toughness measurement in trabecular bone using computed tomography, image correlation and finite element methods. J. Mech. Behav. Biomed. Mater. 109, 1–7 (2020).

46. Allen, M. R. & Burr, D. B. Bone Modeling and Remodeling. in Basic and Applied Bone Biology (2013). doi:10.1016/B978-0-12-416015-6.00004-6

47. Feng, X. & McDonald, J. M. Disorders of bone remodeling. Annu. Rev. Pathol. Mech. Dis. (2011). doi:10.1146/annurev-pathol-011110-130203

48. Van Der Linden, J. C., Homminga, J., Verhaar, J. A. N. & Weinans, H. Mechanical consequences of bone loss in cancellous bone. J. Bone Miner. Res. (2001). doi:10.1359/jbmr.2001.16.3.457

49. Reilly, G. C. & Currey, J. D. The effects of damage and microcracking on the impact strength of bone. J. Biomech. (2000). doi:10.1016/S0021-9290(99)00167-0

50. Yeh, O. C. & Keaveny, T. M. Relative roles of microdamage and microfracture in the mechanical behavior of trabecular bone. J. Orthop. Res. (2001). doi:10.1016/S0736-0266(01)00053-5

51. Oers, R. F. M. van. Simulations of bone remodeling at the cellular scale. Tech. Univ. Endoven 1–114 (2010). doi:10.6100/IR675320

